# Comparison of a novel potentiator of CFTR channel activity to ivacaftor in ameliorating mucostasis caused by cigarette smoke in primary human bronchial airway epithelial cells

**DOI:** 10.1101/2024.03.01.582742

**Authors:** Adrian Constantin Tanjala, Jia Xin Jiang, Paul D.W. Eckford, Mohabir Ramjeesingh, Canhui Li, Ling Jun Huan, Gabrielle Langeveld, Claire Townsend, Daniel V. Paone, Jakob Busch-Petersen, Roman Pekhletski, LiPing Tang, Vamsee Raju, Steven M. Rowe, Christine E. Bear

## Abstract

**Background:** Cystic Fibrosis causing mutations in the gene *CFTR*, reduce the activity of the CFTR channel protein, and leads to mucus aggregation, airway obstruction and poor lung function. A role for CFTR in the pathogenesis of other muco-obstructive airway diseases such as Chronic Obstructive Pulmonary Disease (COPD) has been well established. The CFTR modulatory compound, Ivacaftor (VX-770), potentiates channel activity of CFTR and certain CF-causing mutations and has been shown to ameliorate mucus obstruction and improve lung function in people harbouring these CF-causing mutations. A pilot trial of Ivacaftor supported its potential efficacy for the treatment of mucus obstruction in COPD. These findings prompted the search for CFTR potentiators that are more effective in ameliorating cigarette-smoke (CS) induced mucostasis.

**Methods:** A novel small molecule potentiator (SK-POT1), previously identified in CFTR binding studies, was tested for its activity in augmenting CFTR channel activity using patch clamp electrophysiology in HEK-293 cells, a fluorescence-based assay of membrane potential in Calu-3 cells and in Ussing chamber studies of primary bronchial epithelial cultures. Addition of cigarette smoke extract (CSE) to the solutions bathing the apical surface of Calu-3 cells and primary bronchial airway cultures was used to model COPD. Confocal studies of the velocity of fluorescent microsphere movement on the apical surface of CSE exposed airway epithelial cultures, were used to assess the effect of potentiators on CFTR-mediated mucociliary movement.

**Results:** We showed that SK-POT1, like VX-770, was effective in augmenting the cyclic AMP-dependent channel activity of CFTR. SK-POT-1 enhanced CFTR channel activity in airway epithelial cells previously exposed to CSE and ameliorated mucostasis on the surface of primary airway cultures.

**Conclusion:** Together, this evidence supports the further development of SK-POT1 as an intervention in the treatment of COPD.

## Introduction

Effective therapeutic interventions targeting chronic obstructive pulmonary disease (COPD) remain a significant unmet medical need. In 2019, more than 210 million cases of COPD were reported globally, with COPD accounting for 3.3 million deaths (1). The current standard of care for COPD mostly addresses symptoms and not the underlying pathogenic mechanisms that could reverse the natural course of this condition. Despite some of these interventions, the disease can be progressive, rendering patients vulnerable to lung infections and exacerbations. Mucus aggregation and airway obstruction, early features in COPD-related chronic bronchitis phenotype associated with worse symptoms, as well as accelerated lung function decline, remain largely untreated by current therapies.

Loss of function of the CFTR chloride channel on the surface of airway epithelial cells is known to cause mucus obstruction in the genetic disease, Cystic Fibrosis (2, 3). The environmental factors that induce COPD and bronchitis, including cigarette smoke, reduce CFTR function on the surface of bronchial and tracheal epithelial cultures (4). Together, these findings support the hypothesis that COPD is a form of “acquired” Cystic Fibrosis (5, 6).

Small molecule modulators that augment the function of the CFTR chloride channel, called potentiators, have been shown to ameliorate mucus obstruction and improve lung function in CF patients (7). Ivacaftor (also referred to as VX-770), developed by Vertex Pharmaceuticals, is a highly effective CFTR potentiator approved for the treatment of lung disease in Cystic Fibrosis in patients with genetic mutations associated with decreased CFTR channel function (8). Ivacaftor has been shown to reduce mucostasis and aggregation on the surface of primary, bronchial epithelial cultures exposed to cigarette smoke, the most common in-vitro model of COPD and bronchitis (9). Further, a short, 2 week-long pilot study of ivacaftor in COPD patients revealed modest, non-statistically significant improvements in sweat chloride, nasal potential difference, and symptom scores compared with the placebo. Most recently, Icenticaftor, another oral investigational compound that potentiates CFTR in-vitro (10), was evaluated for 12 weeks in Phase IIb clinical trials in >900 patients with COPD. While there was no significant improvement in lung function measured as the change in through FEV1 from baseline, significant improvement was detected in other symptoms by E-RS cough and sputum scores (11, 12). Altogether, these findings support the hypothesis that potentiators of CFTR channel activity have the potential to reduce mucus obstruction, ameliorate respiratory symptoms and potentially improve lung function.

Both ivacaftor and icenticaftor were discovered in phenotypic screens of compounds that augment forskolin-dependent activation of CFTR channel activity (8, 10). We employed a binding assay using purified Wt-CFTR as the receptor. Using this approach, we discovered a novel small molecule ligand that could bind to either protein receptor and possesses unique structural elements. After conducting chemical modification and screening for ability to enhance Wt and F508del-CFTR channel activity (after temperature rescue), we identified the most potent and efficacious CFTR potentiators. This class of compounds, for which SK-POT is representative, harbours structural elements not shared by Ivacaftor or icenticaftor. In this study, we sought to determine its efficacy in ameliorating mucostasis in human bronchial epithelial cultures exposed to cigarette smoke extract.

## Materials

### Calu-3 cell line

This cell line was obtained from ATCC and tested for mycoplasma contamination at the time of thawing and expansion for the current studies.

### Primary Human Bronchial Cultures

Airway epithelial cultures differentiated at air: liquid interface were derived from non-smoker lung donors of variable ages and provided by the NIH Tissue Culture Facility (University of Iowa) or the Cell Model and Evaluative Core (University of Alabama at Birmingham).

### Ferret tissues

Tracheal explants from wild type ferrets were obtained via overnight shipment from Marshall Farms (NY) and were mounted with intact basement membrane in Ussing chambers for electrophysiologic analysis.

## Methods

### Cell-attached patch clamp studies of CFTR channel potentiation by novel potentiator compound

We used HEK-293 cells stably expressing wild-type human CFTR in patch clamp studies of single channel activity. Cells were the generous gift of D.Rotin (SickKids Hospital, Toronto). These cells were cultured and used as described previously (13). CFTR Cl^−^ channels were recorded in cell-attached membrane patches using an Axopatch 200A patch clamp amplifier and pCLAMP software (both from Molecular Devices, Sunnyvale, CA) (14). The pipette and bath (extracellular) solutions contained 140 mm *N*-methyl-d-glucamine, 140 mm aspartic acid, 5 mm CaCl_2_, 2 mm MgSO_4_, and 10 mm TES, adjusted to pH 7.3 with Tris ([Cl^−^]: 10 mM). The bath was maintained at room temperature.

In order to activate Wt-CFTR channels, 10 uM forskolin was added to the bath. CFTR mediated channel openings were inhibited by application of CFTRInh-172 to the bath. CFTR Cl^−^ channels were potentiated by the addition of SK-POT2 (1 μM) to the bath. To determine channel number, we used the maximum number of simultaneous channel openings observed during the experimental procedure. Single channel recordings were filtered and digitized data as described previously (14). To measure single-channel current amplitude, Gaussian distributions were fitted to current amplitude histograms. To measure *P*_o_, lists of open and closed times were created using a half-amplitude crossing criterion for event detection, and dwell time histograms were constructed and fitted as described previously (14). For the purpose of illustration, single-channel records were filtered at 500 Hz. The resulting Po values and single-channel kinetic parameters were compared with paired *t*-test (Graphpad: Prism); *P* < 0.05 is considered statistically significant.

### Ion transport studies

P2300 Ussing chambers (Physiologic Instruments) were used to measure ion transport across primary human airway cultures (15). CFTR-dependent I_SC_ was assessed by sequential addition of amiloride (100 μM), a chloride gradient plus amiloride (100 μM), forskolin (100 nM), -/+ SK-POT1 and CFTR_Inh_-172 (10 μM) (CFFT, U.S.).

### Studies of the effect of potentiators of CFTR channel activity in CSE-exposed Calu-3

Calu-3 cells were cultured using EMEM (Wisent) supplemented with FBS at 20% (v/v) and 1x Penicillin/Streptomycin (Wisent). Calu-3 cells were seeded on clear-bottom, black-walled 96 well plates (Costar, Corning) at a density of 10,000 cells/well and cultured for 2 days post-confluence. Cells then received a treatment consisting of either cigarette smoke extract (CSE) prepared in 100% DMSO (University of Alabama at Birmingham), dissolved at a concentration of 2% (v/v) in Calu-3 medium, with an equivalent concentration of DMSO alone serving as the vehicle control. The drug treatment was co-applied with CSE, which included SK-POT1 (prepared in 100% DMSO) dissolved in Calu-3 medium at a final concentration of 1 µM. SK-POT1 solutions were prepared from 1 mM stocks. The control for the drug treatment was the vehicle, DMSO at an equivalent volume.

Cells were cultured for an additional 24 hours after chronic treatment application, after which cells were incubated with the proprietary fluorescent membrane potential (FMP) assay buffer. The FMP assay buffer consisted of blue dye (Molecular Devices) dissolved at a concentration of 0.5 mg/mL in chloride-free buffer (150 mM NMDG, 150 mM gluconolactone, 3 mM potassium gluconate, 300 mOsm and pH 7.38). Buffer was incubated with cells at 37 degrees and 5% CO_2_ for 35 minutes. CFTR function was assessed by measuring changes in fluorescence activity after acute CFTR channel activation by addition of cAMP agonist forskolin dissolved in chloride-free buffer to a final concentration 1 µm, using the SpectraMax i3x multimodal plate reader. Fluorescence readings (excitation 530 nm; emission 560 nm) for each well were taken at 30-second intervals for 5 minutes (baseline) or 10 minutes (activation). CFTR activity was then terminated by addition of CFTR-inhibitor 172 dissolved in chloride-free buffer to a concentration of 10 µm, to further verify the specificity of the response to CFTR activity. Fluorescence changes were recorded at 30-second intervals for another 10 minutes. In analysis, all fluorescence values were normalized per well to the final reading prior to the addition of forskolin and expressed as a percentage of this reading.

### Bead Tracking Assay of Mucociliary Movement (MCM)

Differentiated bronchial epithelial cultures, seeded on 24-well transwell inserts were cultured at air-liquid interface. We employed our previously described methods to quantify the velocity of bead tracking, as a surrogate of MCM (16). Ahead of treatment, cultures were incubated with 10 µM N-acetylcysteine dissolved in HBSS (Wisent) for 30 minutes, followed by a final HBSS wash as described previously to reduce the viscosity of the mucous layer. The treatment consisted of either cigarette smoke extract (CSE) prepared in 100% DMSO dissolved at a concentration of 2% (v/v) in UltraG media (the media employed in originally generating the cultures) or DMSO to serve as a control. The potentiators were co-applied with the CSE (or DMSO). Both compounds were prepared in 100% DMSO and dissolved in Ultra G medium at a final concentration of 1 µm. Bronchial epithelia were incubated with the chronic treatment, applied apically, for 24 hours at 37 degrees C and 5% CO2. One hour prior to video imaging, 50 µl of a fluorescent probe suspension in HBSS, consisting of FluoSpheres Polystyrene Microspheres 1.0µm, Green fluorescent or red fluorescent (ThermoFisher Scientific) at 0.02% were placed onto the apical surface of the inserts.

Bead transport rates over the surface of the planar cultures were measured from at 10x and 20x magnifications using an inverted epifluorescence microscope at 37°C with 5% CO_2_ (Nikon eclipse TE2000) with the Hamamatsu orca ER camera capable of taking images at a rate of 9-12 frames per second. A baseline 5-second recording of bead movement was conducted for each cell monolayer prior to apical administration of a 2µl test solution containing vehicle control DMSO, 10 µM forskolin (FSK), or 10 µM FSK with 1 µM VX-770 or SK-POT1. Thirty minutes after addition of test solution, 10x and 20x magnification 5-second videos were repeated. 5 videos were taken per magnification-in the center of the insert as well as in the north, south, east, and west quadrants.; Z-position was adjusted ahead of each video to ensure beads in the airway surface liquid stratum immediately above the cilia were being captured, and to exclude the mostly immobile beads caught in mucus strata further above the cilia.

Upon completion of each set of MCM studies, bead velocity was determined as previously described (16). Bead displacement was determined as the sum of frame-to-frame displacement of each tracked bead. The following formulas were used to generate the tracked displacement (µm) and to determine the velocity of the beads. Velocity (µm/s) = displacement (µm) / 106 (the number of frames the beads were tracked for) x (camera speed in frames per second). The fluorescent beads of each video were enhanced via the contrast enhancement tool and exported as TIFF files in Volocity 6.3. The tracking software, Arivis vision4D, was used to analyze each individual bead velocity and speed using a predetermined tracking pipeline, and the data was exported as an Excel file. The following formulas were used to generate the tracked displacement (µm) and to determine the velocity of the beads. |Velocity| (µm/s) = displacement (m) x 106 x (the number of frames the beads were observed in)-1 x (camera speed in frames per second) (16).

### Statistics

We employed Graphpad Prism (ver.10) for statistical analyses. Each of the dots represent a single biological replicate (cell line plating for Figure 1, 4 and supplementary figure 1) or primary cultures for figures (3) supplementary figure 2). For Figure 5B, each dot represented the mean of 4-5 replicate cultures from individual donors. Paired t-tests were applied for analysis in data in figures 3, 4 and 5b,i. One-way ANOVA with multiple comparison post-test (Tukey) was applied for analysis of data in figure 5b,ii.

**Figure 1.**
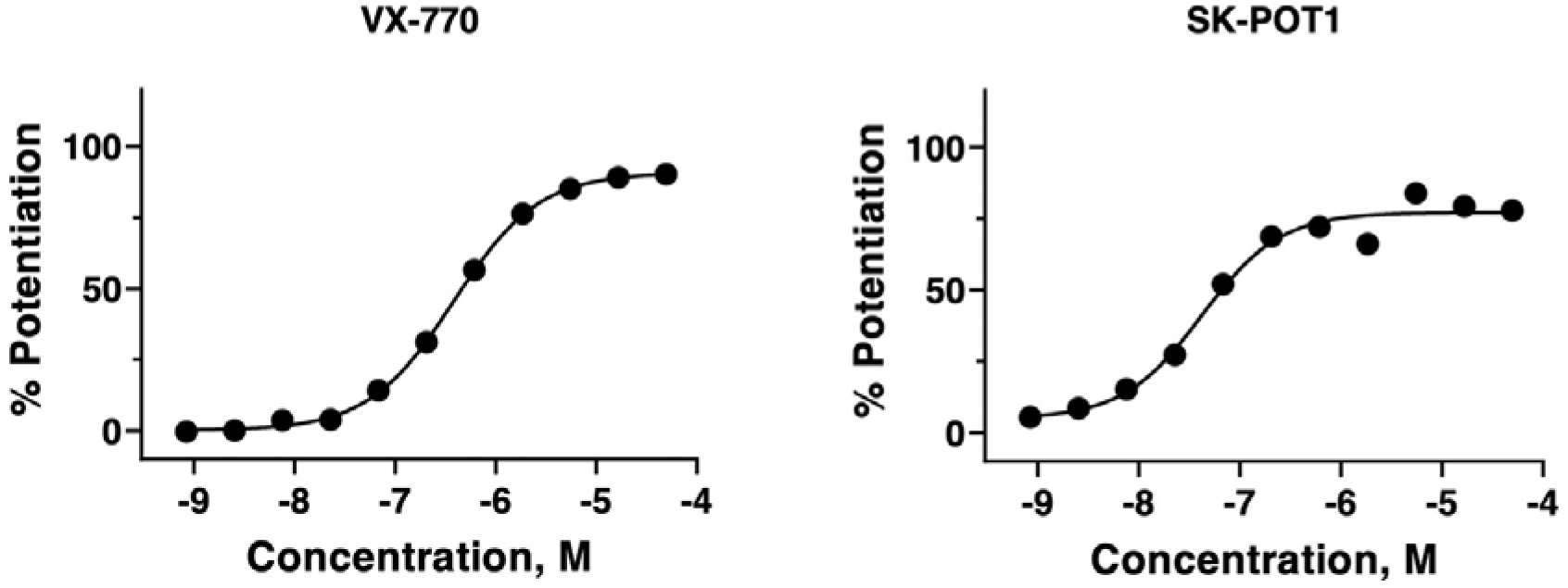
Representative dose responses for VX-770 and SK-POT1 in potentiating cyclic AMP (forskolin activated) chloride channel activity by F508del-CFTR were measured in HEK-293 cells at 30 LJ using the fluorescence based, membrane potential assay (FMP). EC50 for VX-770 is 380 nM and max response (curve fit) is 91% relative to responses to genistein. For SK-POT1, EC50 is 42 nM and max response (curve fit) is 77%. All data were normalized to the response elicited by 10 micromolar genistein plus 30 nanomolar forskolin.

## Results

### A novel potentiator of Wt-CFTR channel activity

A novel class of potentiator molecule was discovered on the basis of its’ affinity to purified and immobilized Wt-CFTR protein and the major Cystic Fibrosis causing mutant protein, F508del-CFTR. Our methods for purifying Wt-CFTR and F508del-CFTR have been published (17–20). After purification, these proteins remained immobilized on Ni-NTA agarose beads. The proprietary technologies for applying and identifying chemical scaffold interactors in the GSK owned DNA-encoded library, has also been described (21). A novel scaffold was identified from the small molecule interactors capable of binding both Wt and F508del CFTR and a series of analogs harbouring the core scaffold were tested for potentiator activity in HEK-293 cells expressing F508del-CFTR. In these studies, cells were cultured at low temperature to ameliorate the mutant’s defective trafficking (22, 23), thereby enabling electrophysiological studies of F508del-CFTR in the plasma membrane. Figure 1, shows a representative dose response of the most potent of the small molecule series derived from the scaffold, i.e., SK-POT1. The fold increase in cyclic AMP dependent F508del-CFTR mediated chloride channel activity induced by potentiator treatment was determined using the fluorescence-based, membrane potential assay previously described (24–26). Analyses of the dose-responses revealed that the mean EC50 (VX-770) was 384 nM, -/+ 86 (SEM) and the maximum responses, 104% of genistein-mediated potentiation for n=4 different cell platings and n=16 dose response studies. Genistein is the first potentiator molecule described for CFTR (27). Interestingly, the mean EC50 for SK-POT1 was 72 nM -/+ 21 (SEM) with a maximum response of 89% of genistein-mediated potentiation. Mean data were derived from 6 independent experiments/cell platings.

Single channel, patch-clamp recordings of cyclic-AMP regulated, channel activity of WT-CFTR expressed in HEK-293 cells are shown in Figure 2. In figure 2,i., we show recordings obtained in the cell-attached mode, where the cells were sequentially exposed to forskolin, then a structural analog of SK-POT1 derived from the same scaffold, followed by the CFTR channel inhibitor, CFTRInh172. While only one channel was activated in the presence of forskolin, a total of 8 channels were stimulated in the presence of the potentiator compound. After the addition of CFTRinh-172, there was a significant decrease in the number of active channels. The bar graph in figure 2,ii, shows the CFTR channel activity after normalization for the total number of channels (n=8). These and replicate studies confirm that this compound potentiates the cAMP-dependent channel activity of Wt-CFTR.

**Figure 2.**
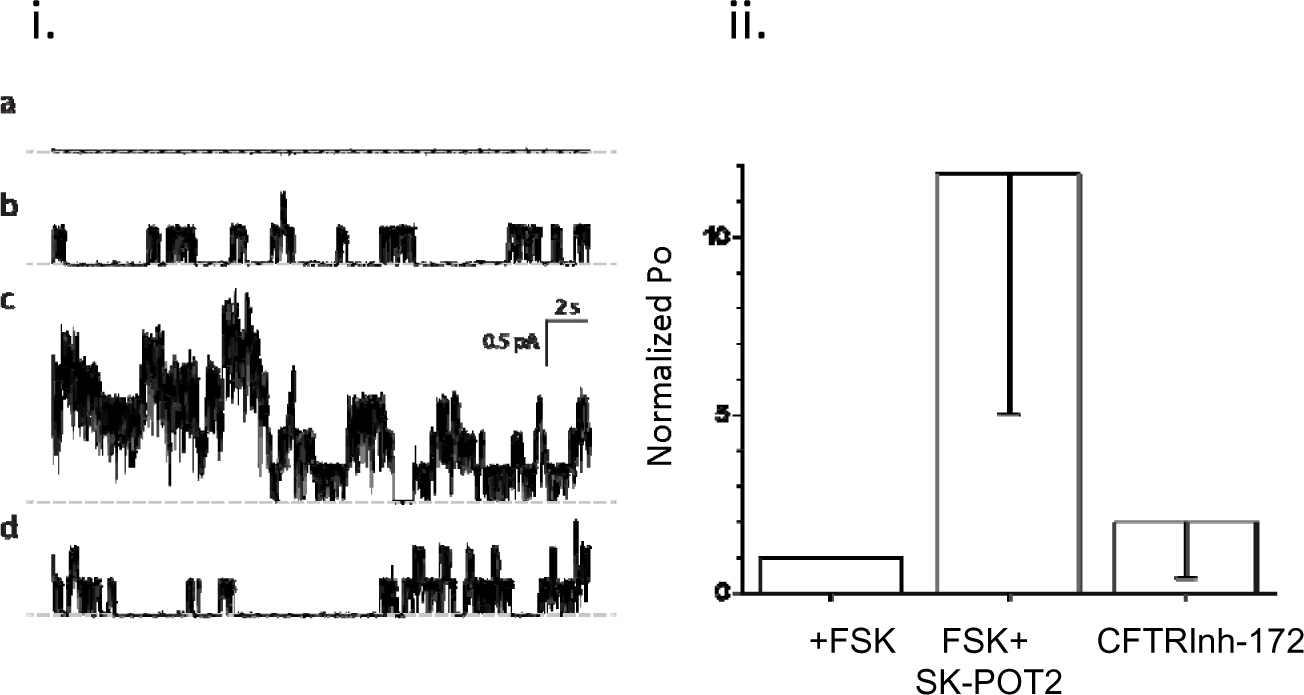
**i. Cell-attached CFTR recordings**. a– representative traces of recordings made in 150 NMDG-Cl base solution complemented with: b – 1 µM forskolin, c – 10 nM SK-POT2 (a structural analog of SK-POT1) and d -10 µM Inhibitor 172; 0 current line represented by gray dashed line; the total number of CFTR channels in the cell-attached patch, which was used for representative traces shown, was estimated as 8. V_m_= -80 mV. ii. Bar graph shows channel open probabilities normalized for total channel number in the presence of forskolin, forskolin plus SK-POT2 or following the addition of CFTR Inh-172. The error bars are standard deviations; n=5 studies for each drug. e – open probabilities for each drug normalized to open probability in presence of 1 µM forskolin (see Methods).

We then tested the impact of the SK-POT1 in potentiating Wt-CFTR in primary human bronchial epithelial cultures. As shown in Figure 3, we show that forskolin activated, short circuit currents mediated by Wt-CFTR in primary bronchial epithelial cultures in Ussing chamber studies, were potentiated by SK-POT1 (1 µM). Representative traces are shown in the left panel and the scatter plots on the right show that these results are reproducible amongst individual donors (shown as individual dots) and statistically significant.

**Figure 3.**
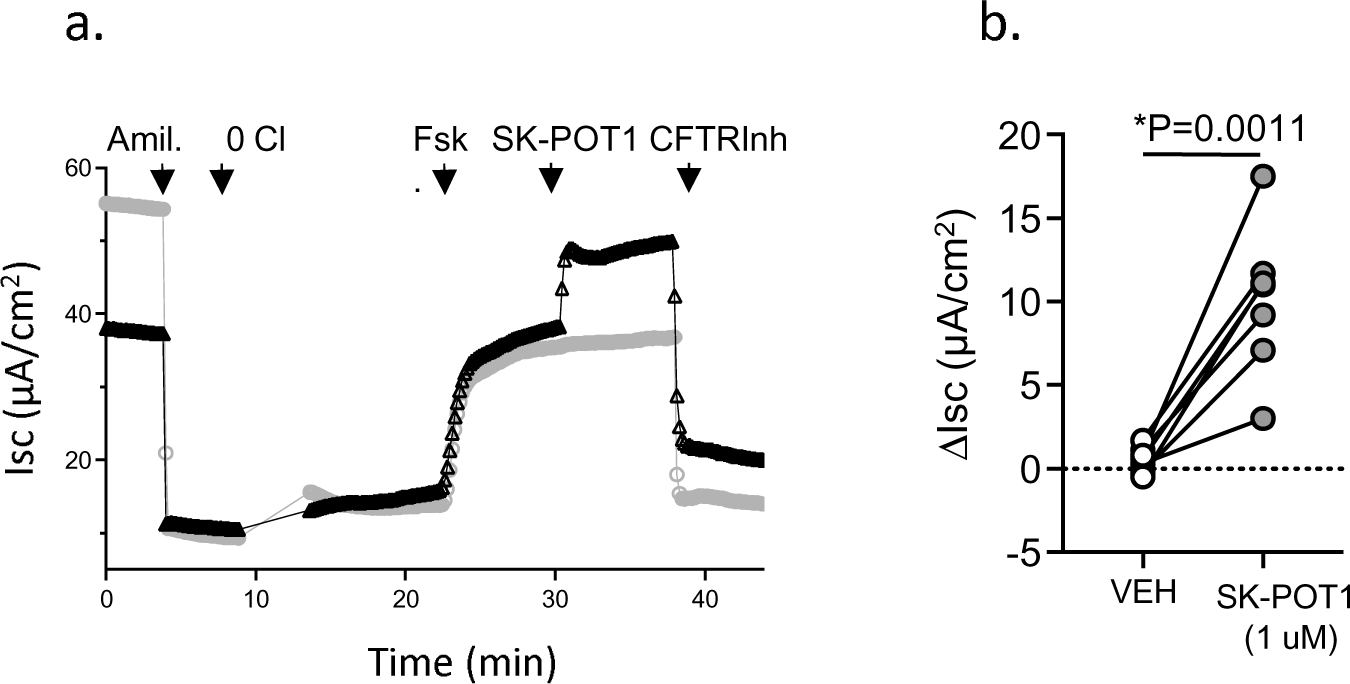
a. Representative Ussing chamber studies of SK-POT1 (1 µM) potentiation after forskolin (FSK) activation (100 nM) in primary bronchial epithelial cultures following amiloride (Amil). b. Dot plot shows 7 paired studies of forskolin-dependent changes in CFTR mediated short circuit with the addition of Vehicle (DMSO) or SK-POT in individual bronchial cultures from 2 donors. The statistical difference between the two interventions was analyzed using the paired “t” test.

We were then prompted to determine if the SK-POT1 compound was capable of ameliorating the negative effect of cigarette smoke (CSE) on Wt-CFTR channel function that was previously reported (9, 28, 29). The cell line, Calu-3, has been employed extensively in studies of the regulation for Wt-CFTR as this airway epithelial cell line endogenously expresses the channel following its differentiation (30). CFTR channel activity was measured using the membrane potential assay (already described earlier - Fig 1) (24, 25, 31, 32). In paired studies, (3 biological replicates, i.e. platings with 3-4 technical replicates, i.e, wells) we assessed the effect of SK-POT1, at 1 uM, in potentiating CFTR activity after CSE exposure. As shown in the bar graph of figure 3, SK-POT treatment led to a significant increase in forskolin-activated CFTR channel activity in the presence of CSE. Representative, time dependent changes in apical chloride conductance in the continuous presence of CSE are shown in Figure 4b. SK-POT1 addition, significantly potentiated forskolin activation of Wt-CFTR in Calu-3 cells.

**Figure 4.**
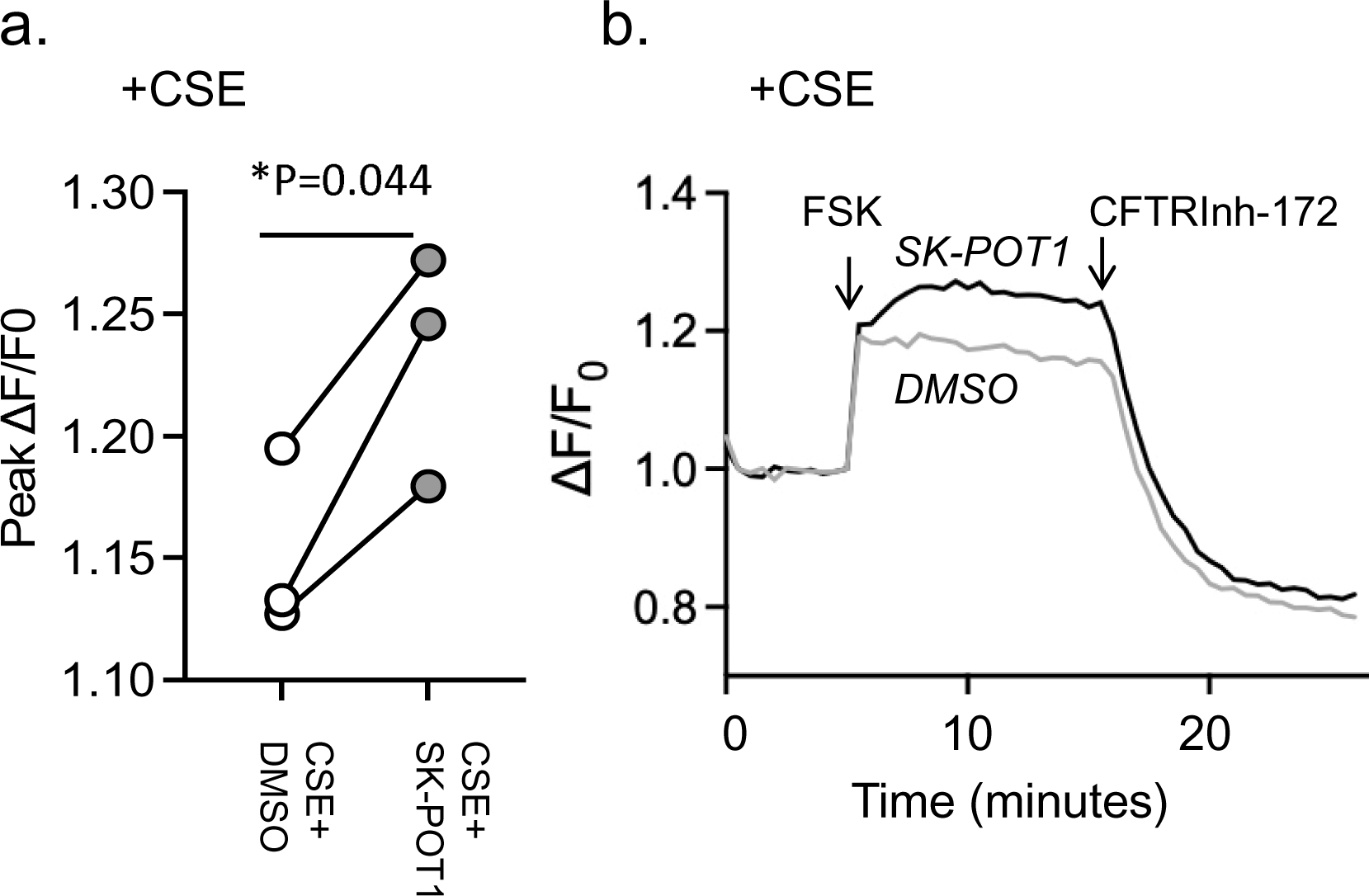
**a**. The scatter gram shows stimulatory effect of SK-POT1 (1 μM) pretreatment on cyclic AMP dependent CFTR channel activity (using the FMP assay) in the presence of cigarette smoke extract (CSE). Each dot pairing corresponds to a separate Calu-3 plating and represents the mean of 3-4 technical replicates, i.e., wells in the plate. The “P”-value was determined using the paired “t” test. b. Left panel: Representative traces showing stimulatory effect of SK-POT1 pretreatment (solid black line) on cyclic-AMP dependent CFTR channel activity in the presence of CSE, relative to DMSO (solid grey line).

Mucostasis, a primary defect in COPD has been modeled in-vitro as reduced movement of fluorescent nanoparticles, seeded in the mucus-containing liquid on top of well-differentiated tracheal airway cultures (16, 33–37). In this and our previously published studies, we seeded fluorescent nanoparticles in the airway surface and tracked bead movement. Bead trajectories are shown in the images of figure 5. The top images (a,i and ii) show that bead displacement is reduced following exposure to CSE as expected (29). We then assessed the effect of pretreatment with the potentiators, VX-770 (1 µM) or SK-POT1 (1 µM) on CSE-altered muco-ciliary movement (panels a.iii-v). In 5 biological replicates (i.e., 5 different donors, 4-5 technical replicates per donor (or videos), we found that pretreatment with SK-POT1 or VX-770 produced a significant increase in bead velocity in CSE treated primary bronchial cultures. Interestingly, the increase in velocity mediated by SK-POT1 exceeded that of VX-770 under the conditions of this study.

**Figure 5.**
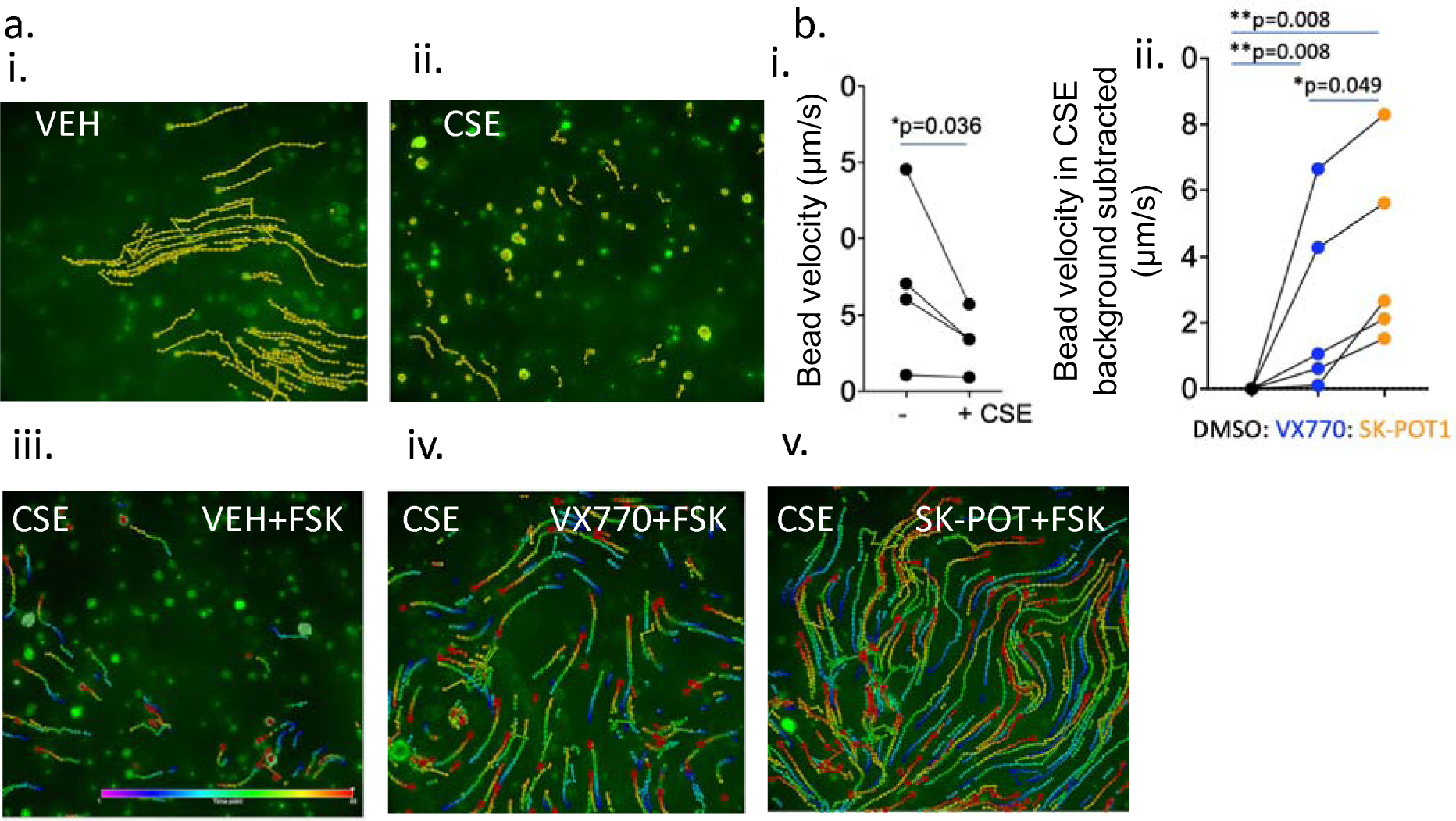
a. i-v, Traces of motile green microspheres over a 5-second period, captured at 10 frames-per-second at 20X objective magnification. Cells were chronically pre-treated for 24 hours with DMSO (i) or 2% cigarette smoke extract (CSE) (ii). CSE-exposed cultures were stimulated with 10 µM forskolin (FSK) plus DMSO (vehicle for potentiators) for 20 minutes at room temperature prior to recording (iii), 1 µM ivacaftor (VX-770) (iv) or 1 µM SK-POT1 (v). Dot colour within the trace represents the point along the 5-second period, with violet (cooler) representing the beginning and red/warmer colours representing the end of the period. b. i, Mean bead velocity for each of 4 donors shown, comparing velocities without CSE to velocities with CSE. ii. Points represent the mean bead velocities for 5 donors and lines compared paired studies. Mean velocities in CSE exposed cultures after VX-770 (blue) or SK-POT1 (orange) pretreatment for each donor normalized to velocity in DMSO. Data in 5.b.i were analyzed using the paired “t” test and data in 5.b.ii analyzed using one way Anova with multiple comparison test (Tukey) of the paired data.

## Discussion

The current studies support the hypothesis that small molecule potentiators of CFTR channel activity have the potential to ameliorate mucus obstruction (38, 39). Further, our findings show that the efficacy of small molecule potentiators to rescue Wt-CFTR channel activity in CSE-exposed airway epithelial cells parallels efficacy in ameliorating CSE associated mucostasis. Further, the novel small molecule potentiator: SK-POT1, is superior to VX-770 in ameliorating the mucostasis induced by CSE in primary airway cultures, at least in the presence of the concentrations employed in this work. Future studies will include comparisons for multiple concentrations of the two potentiators. Altogether, these preclinical studies support the future *in-vivo* evaluation of SK-POT1 as a therapeutic intervention for COPD associated bronchitis.

In contrast to other CFTR potentiators like ivacaftor (VX-770), that were discovered in phenotypic screens of channel activity, the parental molecule from which SK-POT1 was derived, was discovered as a ligand for purified Wt and F508del-CFTR in binding assays. Interestingly, we did not observe a significant difference in the potentiation mediated by VX-770 when SK-POT1 was added simultaneously (Supplementary Figure 1), suggesting that even though they were discovered using different strategies, these two small molecules mediate their activity on CFTR through the same molecular mechanisms.

We observed that the positive effect of the SK-POT1 compound in enhancing CFTR channel activity in CSE exposed epithelial cells, translated to efficacy in rescuing defective mucociliary movement in CSE-exposed primary bronchial cultures (Figure 5). This observation is consistent with the proposal that augmenting CFTR function has the potential to ameliorate mucus aggregation and obstruction in COPD. Interestingly, there was considerable variability in bead velocities (Figure 5), amongst donor-specific cultures. This variability may reflect donor specific differences in the transport properties of primary bronchial cultures as well as certain heterogeneity within each culture, vis a vis the proportion of ciliated cells and mucus secreting cells that can exhibit adherent mucus. While it is not clear how this variability in in-vitro responses will translate to patient-to-patient variance in therapeutic response, several groups have reported concordance between in-vitro responses to CFTR modulators and CF patient specific clinical responses (40–43).

The next steps for these studies will involve preclinical animal studies. The ferret is considered an excellent small animal model for muco-obstructive airway diseases such as Cystic Fibrosis and COPD (15, 44–46). In preparation for a well-powered in-vivo study of SK-POT1 activity in ferrets exposed to cigarette smoke, we first evaluated its activity as a potentiator, in ex-vivo studies of ferret bronchial epithelial cells, cultured at the air-liquid interface using published methods (15) (Supplementary Figure 2). The results of these studies showed that SK-POT enhanced CFTR-mediated, chloride-dependent currents. As evident in the tracing, the imposition of a chloride gradient (basolateral-apical) unmasked basally active CFTR channels, potentially reflecting an altered propensity for endogenous phosphorylation as suggested in a previous study (15). These currents were clearly enhanced by the addition of SK-POT1, a bioelectric phenotype similar to that published for the CFTR potentiator GLPG2196 developed by Galapagos and shown to improve mucus stasis in ferrets (15). In summary, our in-vitro and ex-vivo findings predict a positive in-vivo response to SK-POT1.

## Declarations

### Ethics approval

Studies employing human bronchial epithelial cultures were approved by the University of Iowa Institutional Review Board.

All animal experiments were approved by the Institutional Animal Care and Use Committee of the University of Alabama at Birmingham (IACUC 20232).

### Consent for publication

Not applicable.

### Availability of data and materials

The datasets analyzed during the current study are available from the corresponding author on reasonable request.

---

**Supplementary Figure 1.**
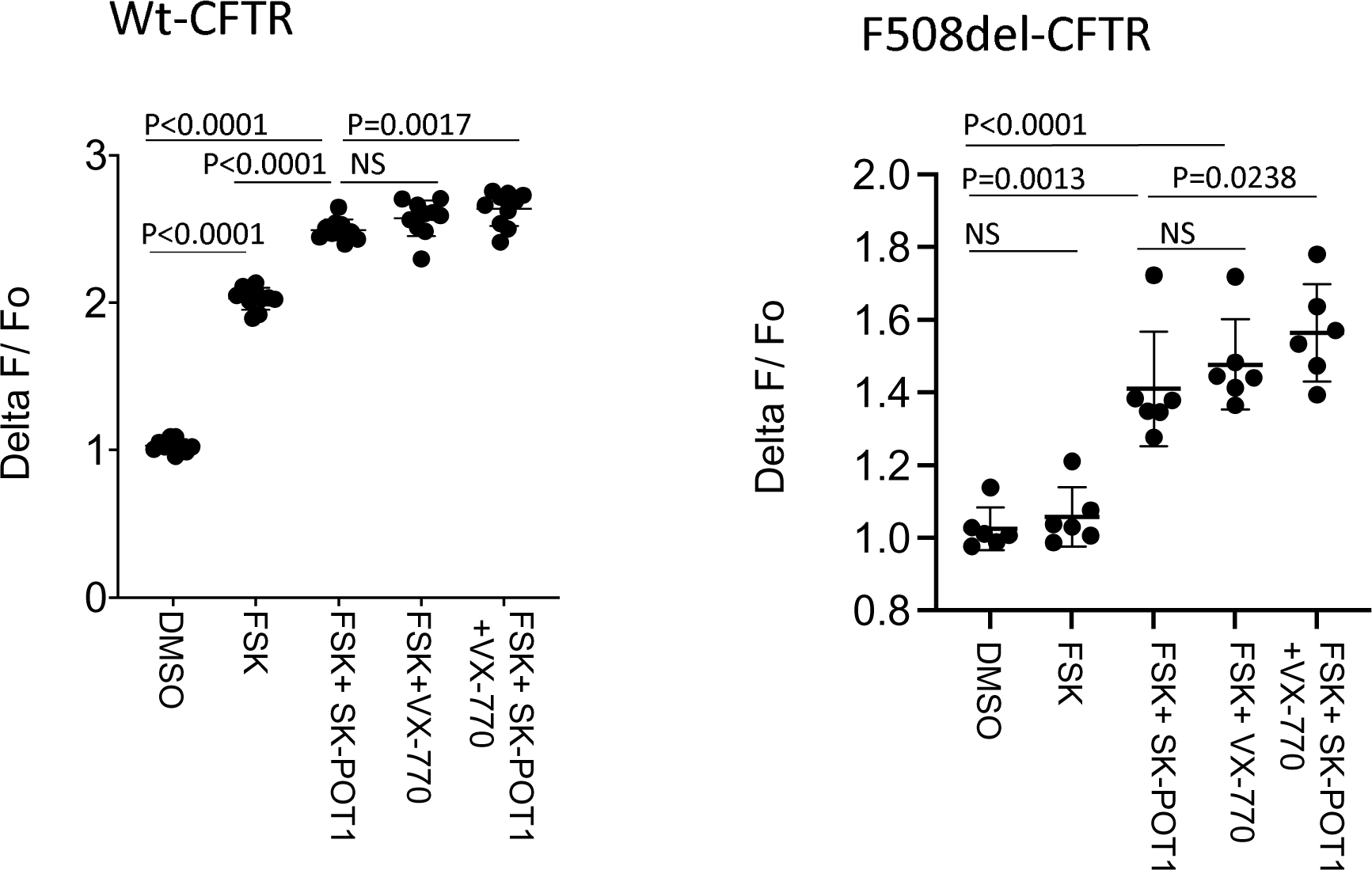
Lack of additive effect of VX-770 (0.1µM) and SK-POT (0.1µM) in potentiating forskolin activated Wt-CFTR or F508del-CFTR (corrected using 3 µM VX-661). Wt and F508del-CFTR were stably expressed in HEK cells. Wt-CFTR was activated using 1 µM forskolin and VX-661 corrected F508del, activated with 10 µM forskolin. For Wt-CFTR and F508del, each symbol reflects a peak response from a sample (well) generated from a total of 3 biological replicates (or cell platings). Mean and SD shown. Mean and SD shown. For Wt and F508del-CFTR, datasets were compared using one-way ANOVA with Tukey’s post-hoc multiple comparison test.

**Supplementary Figure 2.**
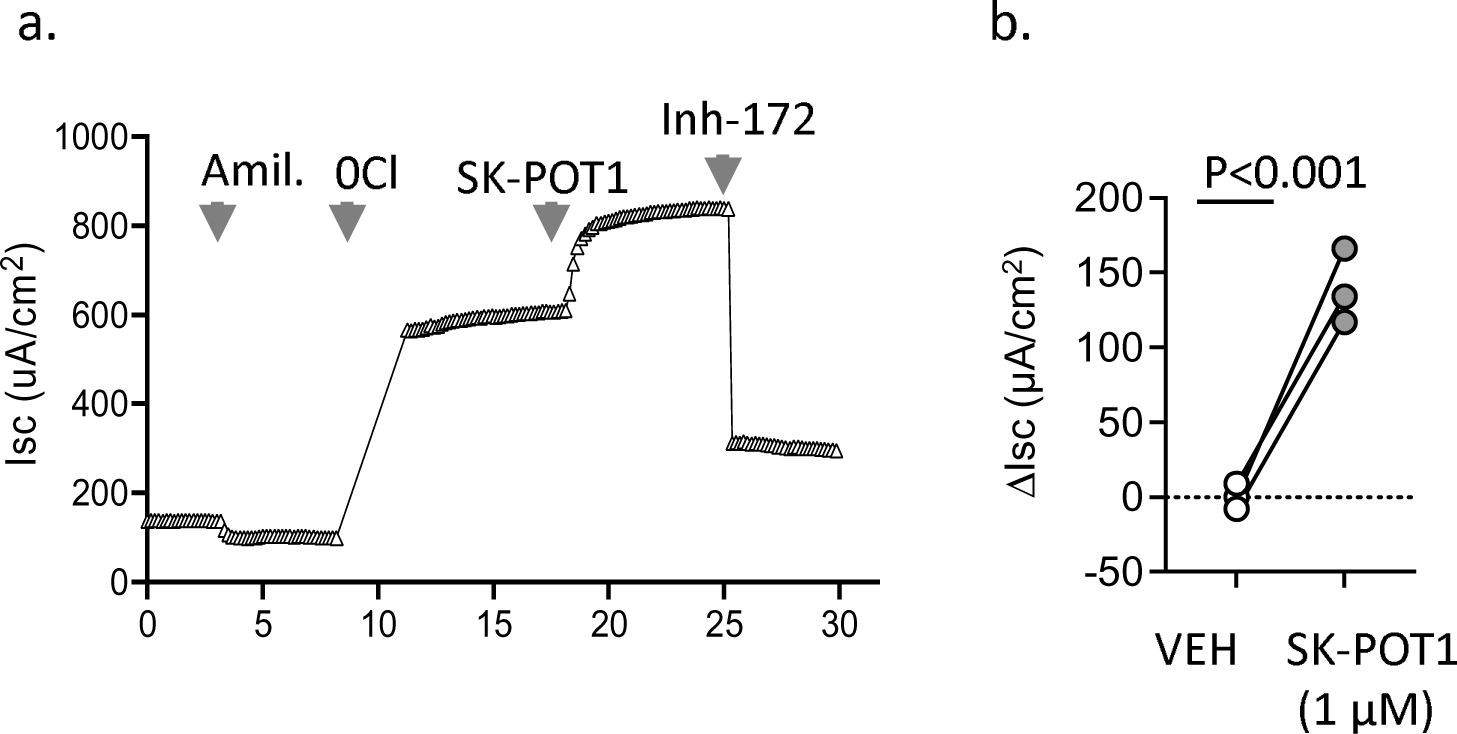
Representative Ussing chamber studies of SK-POT1 (1 µM) potentiation after forskolin activation (100 nM) in primary bronchial epithelial cultures prepared from adult ferret trachea. ii. Scattergram shows forskolin-dependent changes in CFTR mediated short circuit with the addition of SK-POT1 or VEH (n=3 ferret bronchial cultures). Student’s “t” test was conducted.

